# The Human Exonuclease-1 Interactome And Phosphorylation Sites

**DOI:** 10.1101/588574

**Authors:** Wassim Eid, Daniel Hess, Christiane König, Christian Gentili, Stefano Ferrari

**Affiliations:** Institute of Molecular Cancer Research, Winterthurerstrasse 190, CH-8057 Zurich, Switzerland; Department of Biochemistry, Medical Research Institute, University of Alexandria, 21311 Alexandria, Egypt; Friedrich Miescher Institute for Biomedical Research, Maulbeerstrasse 66, CH-4058 Basel, Switzerland

**Keywords:** EXO1, Orbital mass spectrometry, PDCD11, PTMs

## Abstract

Error-free repair of DNA double-strand break is orchestrated by homologous recombination (HR) pathways and requires the concerted action of several factors. Among these, EXO1 and DNA2/BLM execute extensive resection of DNA ends to produce 3’-overhangs, which are key intermediates for downstream steps of HR. To help shedding light on regulatory aspects of DNA repair pathways in which EXO1 participates, we set out to identify proteins interacting with EXO1. Affinity purification of EXO1 followed by Orbitrap mass spectrometry led to the identification of novel partners that are involved in RNA processing or that are the causative agents of rare X-linked disorders. Depletion of a selected subset of EXO1 interacting proteins led to reduction of the DNA damage response. Among those, we examined the RRP5-homologue and NFκB-interacting protein PDCD11/ALG-4, which has roles in apoptosis and is a putative driver gene in cutaneous T-cell lymphoma. We provide evidence that depletion of PDCD11 decreased the formation of γH2AX foci and the phosphorylation of DNA damage response signaling intermediates in response to camptothecin or bleomycin, resulting in increased cellular resistance to DNA damage. Furthermore, extensive coverage of EXO1 sequence (>85%) by mass spectrometry allowed conducting an in-depth analysis of its phosphorylation sites, with the identification of 26 residues that are differentially modified in untreated conditions or upon induction of DNA damage.

As a whole, these results provide the basis for future in-depth studies on novel roles of EXO1 in genome stability and indicate targets for pharmacological inhibition of pathways of cancer development.

**HIGHLIGHTS:**

- Proteome-wide analysis of Exonuclease-1 (EXO1) interacting proteins revealed novel partners involved in RNA processing or that are the causative agents of rare X-linked disorders.
- We provide evidence for a role of PDCD11 in the DNA Damage Response.
- We conducted a comprehensive identification of EXO1 phosphorylation sites.

## INTRODUCTION

The human genome is continuously challenged by different types of insults, with thousands of lesions occurring to DNA every day in each cell [1]. DNA damage arise either as byproduct of normal cellular metabolism and DNA replication or is induced by external factors [2]. DNA double-strand breaks (DSBs) are the most dangerous lesions generated by ionizing radiation (IR), certain chemotherapeutic drugs, collapse of stalled DNA replication forks, endogenous metabolic processes, or arising during meiotic recombination or immunoglobulin diversity generation [3–6]. DSBs are estimated to occur at a rate of ten per cell per day in primary human or mouse fibroblasts [7]. Inappropriate repair of DSBs interferes with DNA replication and transcription and may cause gross chromosomal aberrations [8], resulting in developmental defects, neurodegeneration, aging, immunodeficiency, radiosensitivity and sterility [9] or facilitating the development of cancer through the activation of oncogenes or the inactivation of tumor suppressor genes [10]. DNA damage response (DDR) pathways have evolved as surveillance and protection mechanism to counteract the adverse consequences of DNA lesions and to prevent their transmission to daughter cells [11]. Exonuclease-1 was originally identified in *S. pombe* (Exo1) [12] and subsequently in humans (EXO1) [13] as an enzyme belonging to the Rad2 family of DNA repair nucleases and able to remove mononucleotides from the 5’ end of the DNA duplex [14]. EXO1 is implicated in several DNA repair pathways including mismatch repair, post-replication repair, meiotic and mitotic recombination and double-strand break repair [15–19]. *S. cerevisiae* Exo1 acts redundantly with Rad27 in processing Okazaki fragments during DNA replication [20] and is recruited to stalled replication forks where it counteracts fork reversal [21].

We have previously shown that human and yeast EXO1 are tightly regulated by interaction with CtIP/RBBP8 and 14-3-3 proteins at DSBs and stalled forks, respectively [8, 22, 23]. Additionally, human EXO1 is controlled by post-translational modifications (PTMs), with ATR-dependent phosphorylation targeting it to ubiquitin-mediated degradation upon replication fork stalling [24, 25], and ATM-dependent phosphorylation restraining its activity during homologous recombination [26]. Analogously, Mec1-dependent phosphorylation was shown to inhibit yeast Exo1 activity at uncapped telomeres [27]. Evidence was provided for a role of Cyclin-dependent kinase (CDK)-dependent phosphorylation of human EXO1 in the pathway choice of DSB repair [28]. Adding a further layer of complexity to the regulation of EXO1, we have recently shown that its resection activity is controlled by SUMOylation [29].

Overall, the experimental evidence so far available qualifies EXO1 as a multifaceted protein participating to pathways deputed to the control of genome stability. Considering this, the identification of factors cooperating with EXO1 at sites of damage or controlling its localization and stability in the cell is expected to shed further light on the function of this important nuclease. In this study, we report the identification of novel EXO1 interaction partners. Specifically, we combined Strep-Tactin technology and Orbitrap mass spectrometry (LC-MS-MS) to identify a large set of interacting proteins, among which are ubiquitin E3-ligases, proteins involved in RNA processing and causative agents of rare X-linked disorders. Additionally, the high yield of EXO1 protein allowed identification of novel sites of post-translational modification.

## EXPERIMENTAL PROCEDURES

### Chemicals and antibodies

Camptothecin (Sigma) was dissolved in DMSO at 10 mM concentration. Hydroxyurea (HU) and bleomycin (Calbiochem) were dissolved in water at 1 M and 10 mg/ml concentration, respectively, and filter-sterilized.

Rabbit polyclonal antibodies to EXO1 (F15) and to CtIP were previously described [8, 24]. Additional antibodies used in this study are: rabbit polyclonal to γ-H2AX (9718S, Cell Signaling Technology); rabbit polyclonal and mouse monoclonal to KAP1 (sc-33186 and sc-81411, Santa Cruz Biotechnology), rabbit polyclonal to KAP1-pS_824_ (A300-767A-M, Bethyl), goat polyclonal to ALG4-PDCD11 (sc-163670, Santa Cruz Biotechnology); mouse monoclonal to PHF6 (sc-365237, Santa Cruz Biotechnology); rabbit polyclonal to GFP (ab290, Abcam); mouse monoclonal to GFP (sc-9996, Santa Cruz Biotechnology); mouse monoclonal to MRE11 (12D7, GeneTex); mouse monoclonal to EXO1 (ab4, clone 266, NeoMarkers); rabbit polyclonal to ATM-pS_1981_ (2152-1, Epitomics); mouse monoclonal to MSH6 (clone 44, BD Biosciences) and MSH2 (556349); rabbit polyconal to 14-3-3 (SA-483, Biomol); rabbit polyconal to CHK1-pS345 (2348S, Cell Signaling Technology); rabbit polyclonal to RNF138 (92730, Abcam); GFP-Trap^®^-A (Chromotek).

The anti-rabbit horseradish peroxidase (HRP)-conjugated secondary antibody (A0545) was from Sigma and the HRP-conjugated anti-mouse IgG-k-chain binding protein (sc-51602) as well as the anti-goat HRP-conjugated secondary antibody (sc-2350) were from Santa Cruz Biotechnology.

### Cloning strategy

A fusion construct between human EXO1 and the Strep-Tag was generated by subcloning full-length human *EXO1b* cDNA in the mammalian expression vector pEXPR-IBA3^®^(IBA BioTAGnology), which allows addition of a C-terminal tag to the protein of interest. The coding region of human EXO1b (codons 1-846) was amplified by PCR using pcDNA3.1-HisC-EXO1b [24] as template and the primers (RE sites underlined):

> 5’-ATGGGAGA<u>CCGCGG</u>GATACAGGGATTGCTACAATTTATC-3’ (forward);
>
> 5’-CTCGAG<u>GGATCC</u>CTGGAATATTGCTCTTTGAACACGG-3’ (reverse).

The PCR product was inserted between the *SacII* and *BamHI* restriction sites of pEXPR-IBA3 to generate a translational fusion between EXO1b and a tandem StrepTag, separated by a two-amino acid spacer. The construct was verified by sequencing.

### Cell culture

HEK-293T human embryonic kidney cells and U2-OS human osteosarcoma cells (American Type Culture collection, Manassas, VA, USA) were maintained in Dulbecco’s modified Eagle’s medium (DMEM; Invitrogen) supplemented with 10% fetal calf serum (Gibco) and penicillin/streptomycin (100 U/ml; Gibco).

For transfection experiments, cells were seeded in 10 cm dishes and allowed to adhere overnight. Cells were transiently transfected with constructs of interest or the empty vector using 1 μg of the DNA and 4 μl of the transfecting reagent Metafectene (Biontex, Germany) according to the manufacturer instructions. Cells were collected and lysed 48 h after transfection using protocols for isolation of total proteins [8].

When required, cells were treated with either HU (2 mM, 16h), CPT (1 μM, 4h) or bleomycin (10 μg/ml, 1h). Alternatively, cells were exposed to 10 Gy ionizing radiation using a Faxitron Cabinet X-ray system, model 43855D (Faxitron X-ray Corp., Wheeling, IL, USA), and harvested 1h post-exposure.

### RNA interference

Two oligonucleotides for each target were purchased at Qiagen (Switzerland) and transfected at 40 nM concentration using Truefect-Lipo (United BioSystems Inc. USA) according to the manufacturer instructions. Experiments were typically performed for 48-72 h.

### Cell viability

Cells were seeded and grown for 24h in full medium. For RNAi experiments, cells were transfected with control or specific siRNA oligonucleotides and, after 24h, re-seeded in 96-well plates at density 1-1.5 × 10^3^/well and grown for further 24h. DNA damaging agents were administered at the range of concentrations indicated in the figures. Cell viability was assessed using a Resazurin-based a fluorometric assay (Promocell, Germany) [30] 48h after treatment with DNA damaging agents.

### Proteins pull-down

Upon 48h expression, cells were harvested in ice-cold lysis buffer. Whole cell lysates (~20 mg) from mock or Strep-EXO1 expressing cells were incubated on a wheel rotator for 1h at 4°C with 70 μl of Strep-Tactin^®^ Sepharose beads (IBA BioTAGnology) pre-equilibrated in lysis buffer. Beads were washed 3 times with 1 ml of ice-cold lysis buffer and incubated with 100 μl of 5 mM D-Desthiobiotin (IBA BioTAGnology) – a high-affinity agonist - at 4°C for 2h with occasional shaking. Beads were centrifuged, the supernatant was transferred to a new tube and competition with D-Desthiobiotin (100 μl) was repeated once. Supernatants from the two elution steps were pooled and subjected to tri-chloracetic acid (TCA) precipitation by addition of the acid to a final concentration of 20%. Samples were vortexed, incubated on ice for 2h and centrifuged. Protein pellets were washed in 1 ml of 10% ice-cold TCA, vortexed and centrifuged. Finally, pellets were washed with 1 ml of ice-cold HPLC-grade acetone, centrifuged and air-dried. Protein pellets were stored at −80°C until the time of analysis.

### Silver staining

All reagents were prepared in MilliQ-ddH_2_O and solvents were of HPLC quality. Polyacrylamide gels were fixed in 50% methanol, 5% acetic acid in ddH_2_O for 20 min, washed with 50% methanol in ddH_2_O, then additionally for 10 min in ddH_2_O. The gel was soaked in 0.02% sodium thiosulfate for 1 min, washed twice with ddH_2_O for 1 min, soaked in cold 0.1% silver nitrate solution for 20 min at 4°C in the dark. The gel was then washed twice with ddH_2_O for 1 min and developed in 0.04% formaldehyde in 2% sodium carbonate with continuous, gentle shaking. Development was terminated by addition of a 5% acetic acid solution.

### Mass spectrometry

Silver stained protein bands were excised from the gel, reduced with 10 mM DTT, alkylated with 55 mM iodoacetamide and cleaved with porcine trypsin (Promega, Madison, USA) in 50 mM ammonium bicarbonate (pH 8.0) at 37°C overnight. The extracted peptides were analyzed by capillary liquid chromatography tandem mass spectrometry (LC-MSMS) using a Magic C18 100 μm × 10 cm HPLC column (Swiss BioAnalytics, Switzerland) connected on line to a 4000 Q Trap (MDS Sciex, Concord, Ontario, Canada) as described earlier [31].

TCA precipitated Strep-EXO1 pellets were washed with acetone, reduced with tris(2-carboxyethyl)phosphine (TCEP), alkylated with jodoacetamide and digested with trypsin. Tryptic peptides were injected into a reversed phase column for liquid chromatography-mass spectrometry (LC-MS) analysis in the information-dependent acquisition mode. Electrospray ionization LC-MSMS was performed using a Magic C18 HPLC column (75 μm × 10 cm; Swiss BioAnalytics) with a 1200 Nano-HPLC system (Agilent Technologies) connected to a LTQ Orbitrap Velos (Thermo Scientific). The peptides were loaded onto a peptide cap-trap (Michrom BioResources) at a flow rate of 10 μl/min for 5 min and eluted at a flow rate of 400 nl/min with a linear gradient of 2–36% acetonitrile in 0.1% formic acid and 0.005%TFA in H_2_O in 30 min. Information-dependent acquisition analyses were done according to the manufacturer’s recommendations, i.e. 1 survey scan (MS1) at 60K resolution in the Orbitrap cell was followed by up to 20 product ion scans (MS2) in the linear ion trap, and precursors were excluded for 15 sec after their second occurrence. To improve both sequence coverage and number of identified proteins, the analyses were done 4 times with different mass ranges for MS1 (gas phase extraction). The mass ranges were as follows: 400-1200, 400-600, 600-800 and 800-2000. Individual MS2 spectra, containing sequence information were compared with the program Mascot against the human subset of the protein sequence database Swiss-Prot 2010_09 [32]. Carboxyamidomethylation of cysteine (+57.0245 Da) was set as a fixed modification and oxidation of methionine (+15.9949 Da), phosphorylation (+79.966331 Da) of serine and threonine and acetylation of protein N-termini (42.0106 Da) were set as variable modifications. Parent tolerance was 10 PPM (monoisotopic) and fragment tolerance 0.6 Da (monoisotopic). The MASCOT results were evaluated with the software Scaffold and Scaffold PTM (www.proteomesoftware.com) to further validate the phosphorylation sites. Additionally, peptides were quantified with the software Progenesis LC (www.nonlinear.com).

The mass spectrometry proteomics data have been deposited to the ProteomeXchange Consortium [33] via the PRIDE partner repository, with dataset identifier PXD002780.

### Western blot and immunoprecipitation

Proteins were extracted using ice-cold lysis buffer (50 mM Tris–HCl pH 7.5, 120 mM NaCl, 20 mM NaF, 1 mM EDTA, 6 mM EGTA, 15 mM Na-pyrophosphate, 0.5 mM Na-orthovanadate, 1 mM benzamidine, 0.1 mM phenylmethylsulfonyl fluoride and 1% Nonidet P-40). Protein concentration was determined using the Bio-Rad Protein Assay Reagent (Bio-Rad). Proteins were separated on SDS-polyacrylamide gels, transferred to polyvinylidene difluoride (PVDF) (GE-Healthcare) and probed with appropriate antibodies. Immunoblot analysis was performed using BioRad Clarity™ Western Substrate (#170-5060) and proteins were visualized with the FUSION SOLO^®^ chemiluminescence imaging system (Vilber) [25]. Immunoprecipitation was performed as described previously [24]. To ensure that the observed interactions were not DNA-mediated, ethidium bromide was included in all samples during immunoprecipitation.

### Immunofluorescence microscopy and image analysis

U2-OS cells grown on cover slips were fixed in 4% formaldehyde (w/v) in PBS for 15 min at room temperature (RT). Cover slips were incubated overnight at 4 °C with primary antibodies followed by Alexa–conjugated secondary antibodies for 1h at RT. The cover slips were mounted with Vectashield^®^ (Vector Laboratories) containing DAPI. Images were acquired using a Leica fluorescence microscope DM6-B. Nuclei were detected based on the DAPI signal and, for each nucleus, the intensity of DAPI (blue) and γ-H2AX (red) signals were extracted using the open-source software, Cell Profiler [34]. Data were plotted using GraphPad Prism 8.0.2.

### Gene network analysis

To assign links between EXO1 and cellular processes through its interacting proteins, gene networks were constructed. The literature-based ResNet Mammalian 11.0 Database in Pathway Studio 11.0 (Elsevier, Amsterdam, Netherlands) [35] was used to determine pathway and cellular processes in which EXO1-interacting proteins have an annotated function. Each pathway link is supported by at least one published reference.

## RESULTS

### Identification of EXO1 interacting proteins

To the end of identifying EXO1 interacting proteins, we created a fusion construct encoding EXO1 and a tandem Strep-tag (EXO1-2x-Strep, EXO1-Strep hereafter). EXO1-Strep was expressed to an extremely low level (Fig. 1A, lane 2), in a manner reflecting the amount of the endogenous protein [24], and became clearly detectable upon pull-down with Strep-Tactin^®^ beads. The recombinant fusion protein displayed interaction with the established partner CtIP (Fig. 1A, lane 4) [8]. Lack of binding to MRE11 (Fig. 1A, lane 4), which served as negative control [8], confirmed the suitability of the EXO1-Strep construct for protein interaction studies. Next, EXO1-Strep was expressed on a large-scale in HEK293T cells that were treated in the presence or the absence of HU or IR. Approximately 1% of the material deriving from HEK293T cells was examined by Western blot to confirm activation of the DNA damage response by HU or IR (Fig. 1B) and the pulled-down recombinant fusion protein was visualized by silver staining (Fig. 1C). Orbitrap mass spectrometry (ProteomeXchange accession: PXD002780) yielded a number of established (14-3-3ζ, 14-3-3β, 14-3-3ε, MSH6) [22, 23, 36] and novel EXO1-interacting proteins (Table 1), upon subtraction of frequently recurring background contaminants [37]. Among novel interactors, a large proportion of hits displaying a ratio >5 for spectra counts in EXO1-Strep over mock-transfected cells consisted of proteins involved in RNA metabolism. These were components of ribonucleosomes (hnRNP-A, -R, -G, -Q), factors required for mRNA splicing (SFRS7, SFRS5) and regulators of heterochromatin integrity (HP1B3). A subset of the hits displayed links to ubiquitin-dependent pathways, such as GNL3 (a stabilizer of the ubiquitin E3 ligase MDM2), CHIP/STUB1 and PRP19 (ubiquitin E3 ligases). An interesting novel interacting protein with significant spectra counts ratio was the PHD-finger protein 6 (PHF6), a transcriptional regulator whose mutation causes the X-linked Boerjeson-Forssman-Lehmann syndrome (BFLS) (OMIM 301900). Additional interacting proteins of interest displaying a ratio >5 were: the cell proliferation-associated antigen (KI67), a ribosome biogenesis and assembly factor (MRT4), a double-stranded RNA binding protein (STAU1), a serine/threonine phosphatase (PGAM5), ribosomal RNA processing factors (RRP1B, RRP5/PDCD11), a translational repressor (FMR1) whose mutation causes the Fragile X Syndrome (OMIM 300624), and proteins of as yet unknown function (GPTC4, ZNF655).

**Figure 1.**
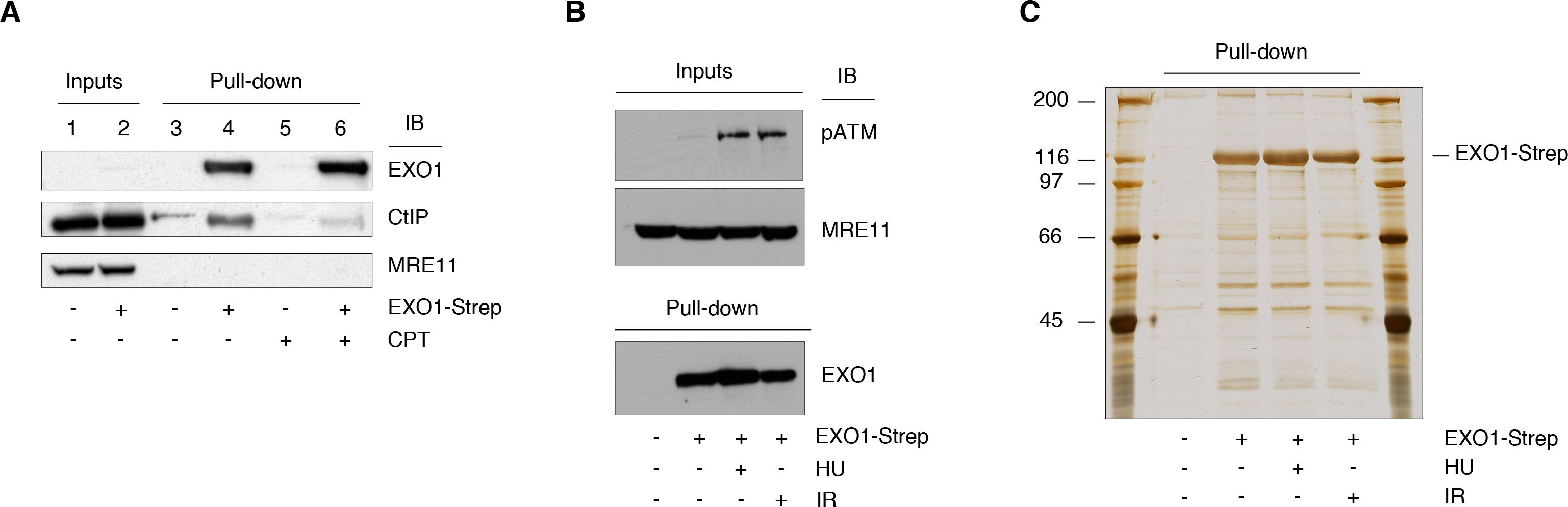
Characterization of EXO1-Strep. A. EXO1-Strep was expressed in HEK293T cells that were either left untreated or treated with camptothecin (CPT, 1 μM) for 4h. EXO1 was pulled down from whole cell extract (WCE, 1 mg). The membrane was probed with the indicated antibodies. B. WCE (50 μg) obtained from mock and EXO1-Strep transfected cells treated with HU or IR were examined by Western blot with the indicated antibodies. MRE11 was used as loading control. C. WCE (18 mg) from the cells shown in panel B underwent pull-down with Strep-Tactin^®^ beads. A fraction of the pulled down material was analyzed by SDS-PAGE and visualized by silver staining.

### Validation and characterization of selected EXO1 interacting proteins

To assess the validity of the mass spectrometry data using an independent method, we conducted immunoprecipitation studies on extracts of HEK-293T cells ectopically expressing GFP-EXO1 (Fig. 2A). Immunoblot analysis confirmed interaction between GFP-EXO1 and PHF6 (Fig. 2B), GFP-EXO1 and RNF138 (Fig. S1), as well as established partners such as MSH2, MSH6 and 14-3-3 proteins that were taken as positive controls (Fig. 2B).

**Figure 2.**
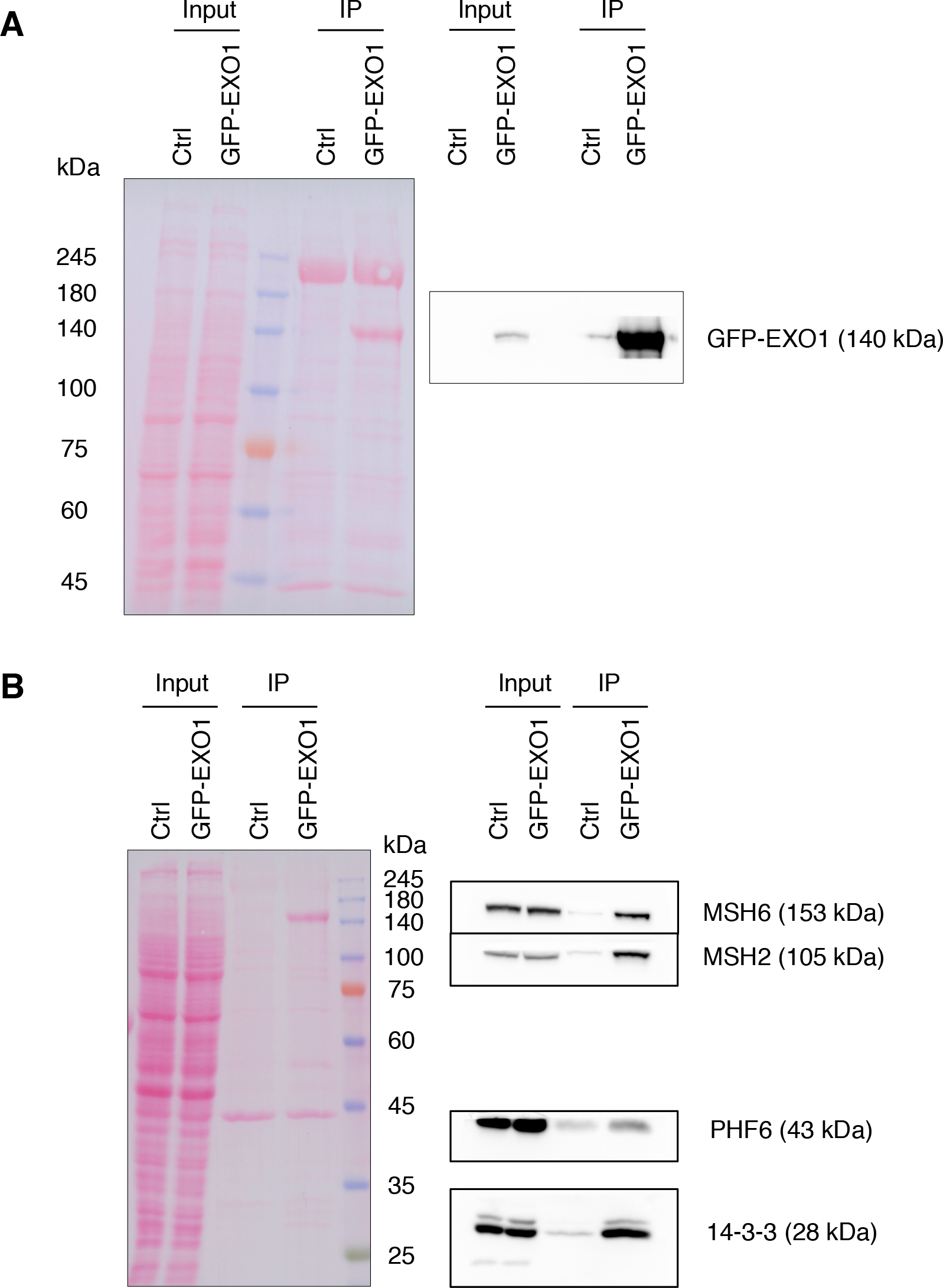
Verification of EXO1 interacting proteins. A. HEK-293T cells were transfected with an empty plasmid or a plasmid expressing GFP-EXO1 and WCE (2 mg) submitted to immunoprecipitation with an antibody to GFP. Upon transferring proteins to PVDF, GFP-EXO1 was revealed with a specific antibody. Inputs correspond to 50 μg WCE. Ponceau-red (PR) stained proteins were used as loading control. B. HEK293T cells were transfected as above were immunoprecipitated with an antibody to GFP and examined with the indicate antibodies. Ponceau-red (PR) stained proteins were used as loading control.

To assess the impact of novel EXO1-interacting proteins on the DNA damage response, we proceeded by knocking down the expression of a selected set of proteins in U2-OS cells. Under these conditions, we quantified γ-H2AX foci formation in response to treatment with the topoisomerase-1 inhibitor camptothecin, a chemotherapeutic agent known to induce DSBs exclusively during DNA replication through the trapping of DNA topoisomerase 1 cleavage complexes [38]. Immunofluorescence analysis showed a reduction of the DDR marker γ-H2AX upon depletion of all selected proteins (Fig. 3 and Fig. S2). Down-regulation of PDCD11 expression appeared particularly effective in decreasing DDR. Since the RRP5-homologue and NFκB-interacting protein PDCD11/ALG-4 has roles in apoptosis [39], in the generation of mature 18S rRNA [40] and its absence is frequently observed in cutaneous T-cell lymphoma [41], we decided to further characterize this candidate protein.

**Figure 3.**
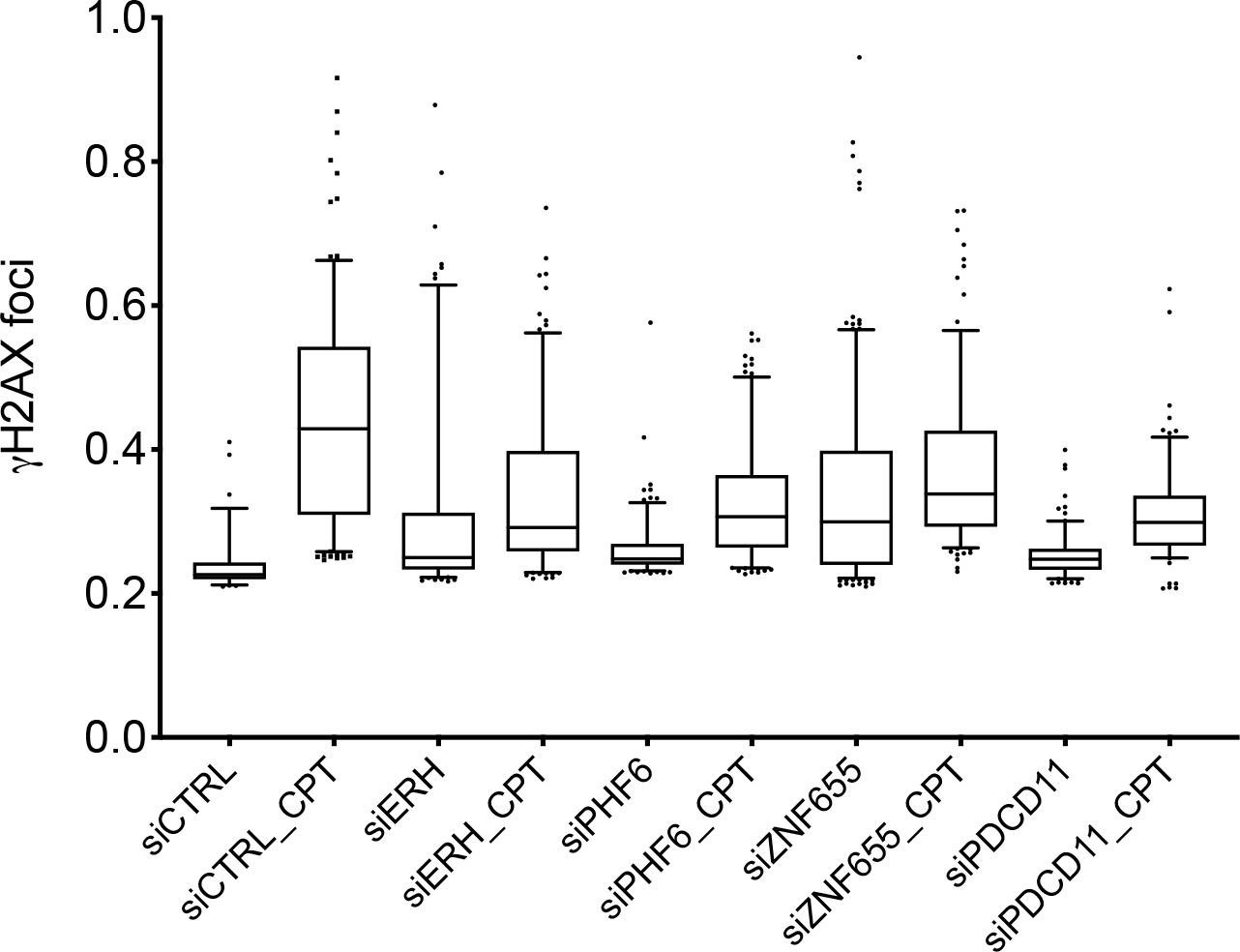
Depletion of EXO1 interacting proteins affects the DNA damage response. U2-OS cells were transfected with the indicated siRNAs (two oligonucleotides per target) and 48h upon transfection treated with camptothecin (CPT, 1 μM) for 4h. Cells were fixed, decorated with an antibody to γ-H2AX and nuclei counterstained with DAPI. DNA damage foci were visualized with a 60x oil-immersion objective. An average of 300 cells for each condition was examined using Cell Profiler. Data were plotted using GraphPad Prism 8.0.2. One of three independent experiments is shown.

We initially assessed the efficiency of the two siRNA oligonucleotides used to deplete PDCD11 expression. The data showed that both siRNAs were effective in diminishing PDCD11 protein level upon 48h transfection of U2-OS cells (Fig. S3A). Cell cycle analysis showed that downregulation of PDCD11 caused a slight increase of the G1 cell population and a parallel decrease of S and G2-M cells (Fig. S3B and S3C). Cell survival assays upon exposure of cells to camptothecin or the IR-mimetic bleomycin revealed that PDCD11 depletion conferred higher resistance to the DNA damaging agents as compared to siRNA control depleted cells (Fig. 4). Analysis of the DDR in distinct cell cycle populations revealed that both in G1 and G2-M cells there was a reduced response to the DNA damaging agent bleomycin (Fig. 5A and 5B). Western blot analysis of the DDR markers CHK1 and KAP1/TRIM28 revealed a decrease of phosphorylation for both proteins in the PDCD11 depleted cells treated with bleomycin (Fig. 5C).

**Figure 4.**
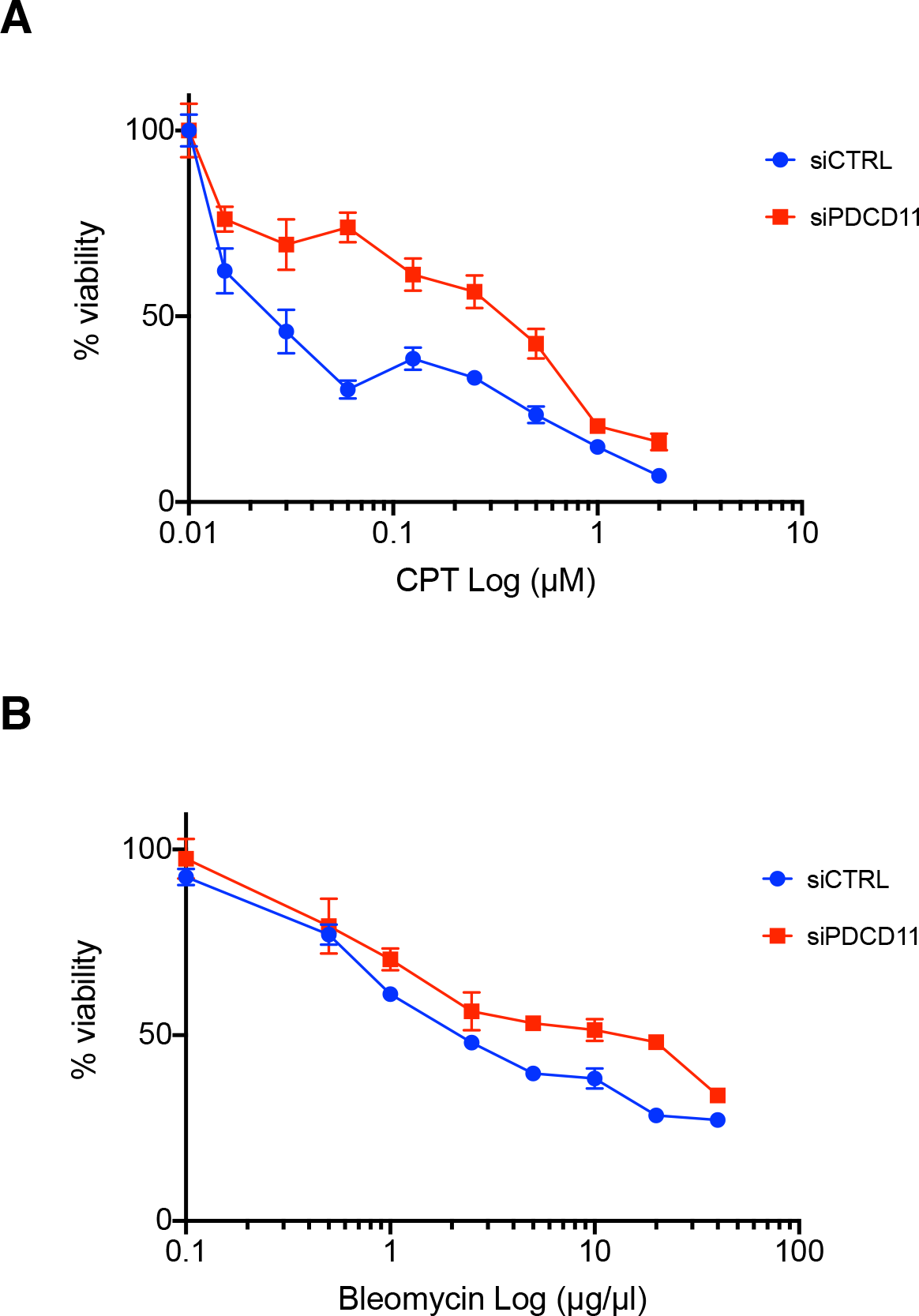
PDCD11 depletion increases resistance to DNA damage. A. U2-OS cells were transfected with CTRL siRNA or a combination of two siRNA oligonucleotides to PDCD11, treated in the presence or the absence of camptothecin and cell viability determined as described in Materials and Methods. One of two independent experiments performed with triplicate determinations is shown. B. U2-OS cells were transfected with CTRL siRNA or a combination of two siRNA oligonucleotides to PDCD11, treated in the presence or the absence of bleomycin and cell viability determined as described in Materials and Methods. One of two independent experiments performed with triplicate determinations is shown.

**Figure 5.**
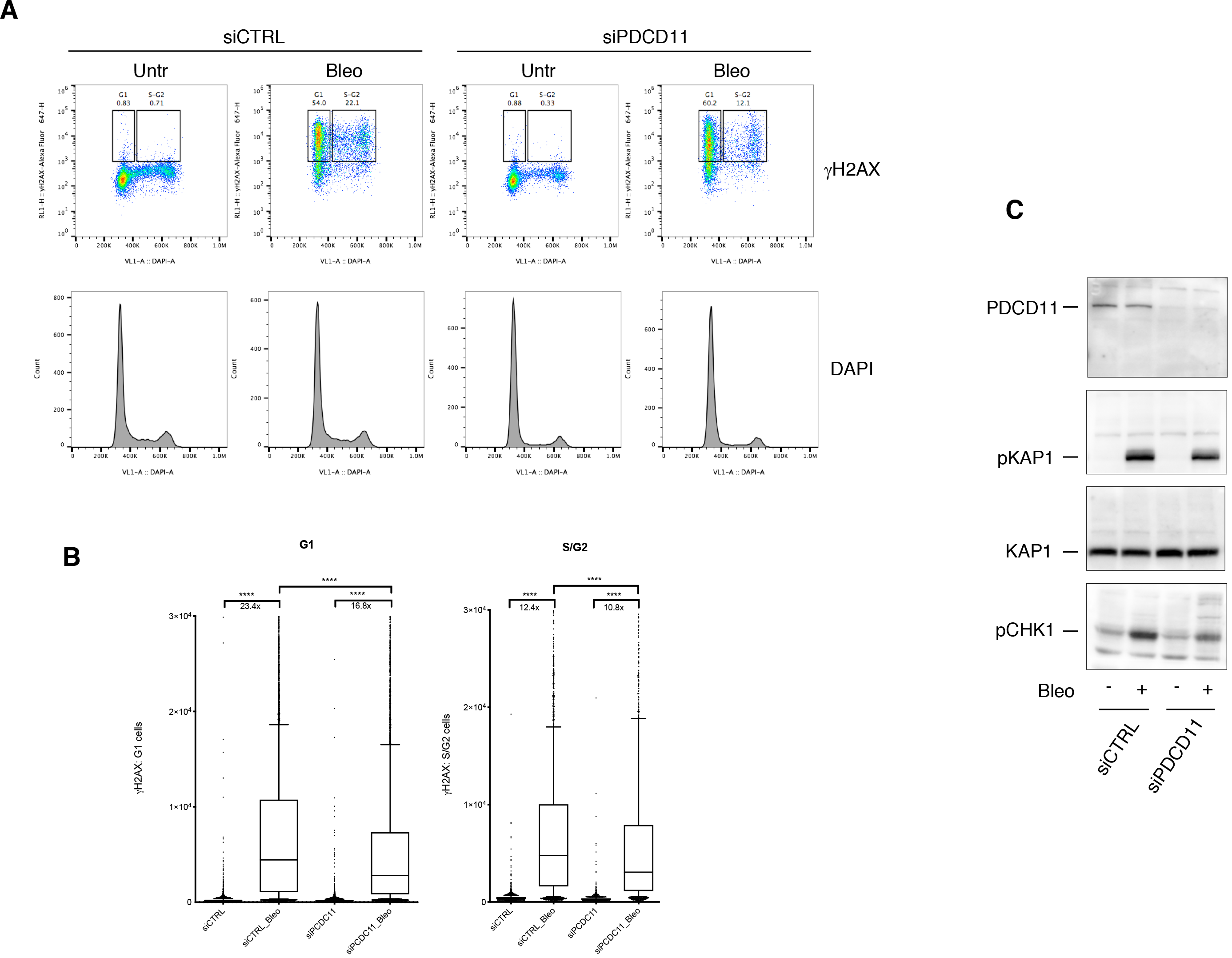
PDCD11 depletion decreases the DNA damage response. A. U2-OS cells were transfected with CTRL siRNA or a combination of two siRNA oligonucleotides to PDCD11. 72h upon transfection cells were treated with bleomycin (1h, 10 μg/ml), fixed, and γH2AX foci were detected by flow cytometry. Dot plots (top) and histograms (bottom) show γH2AX and DAPI signals, respectively. One of two independent experiments is shown. B. Quantification of the events gated in panel A (top). C. Western blot analysis of DDR markers in total extracts of the cells described in panel A. KAP1 was used as loading control.

Taken together, these data indicate that the EXO1 interacting protein PDCD11 participates in the DNA damage response.

### EXO1 sites of phosphorylation

Isolation of Strep-EXO1 from HEK293T cells yielded an amount of protein sufficient to cover 86% of EXO1 sequence by Orbitrap mass spectrometry. Hence, post-translational modifications such as phosphorylation could be reliably scored. The data showed that EXO1 contained 26 sites of phosphorylation in all conditions examined (Table 2). Analysis of peptide features revealed that the majority of the sites that we identified fell into the two largest families of phosphorylation sites described in the literature, namely targets of proline-directed kinases or of acidophylic kinases, accounting in total for about 77% of all detected phosphorylation sites [42]. To the proline-directed kinases class of sites belong residues 475, 598, 621, 623, 639, 814 and 824, whereas to acidophylic kinases target sites belong amino acids 422, 426, 700, 702, 744, 745 and 746 (Table 2). Among the phosphorylated residues that we have previously described in EXO1 [25], this study confirmed the presence of S454 and S714 as DNA damage-dependent sites (Fig. 6). Assessment of peptide intensity upon HU treatment, normalized to the untreated condition, showed an increase of phosphorylation at S454, at the novel target double-site T621/S623 and particularly at the established S714 [25, 26] (Fig. 6). Less prominent increase of phosphorylation under these conditions was observed at S422 and S426 (Fig. 6). On the other hand, IR appeared to particularly increase phosphorylation at S385 and, to a lesser extent, S714 (Fig. 6), in line with our previous reports [25].

**Figure 6.**
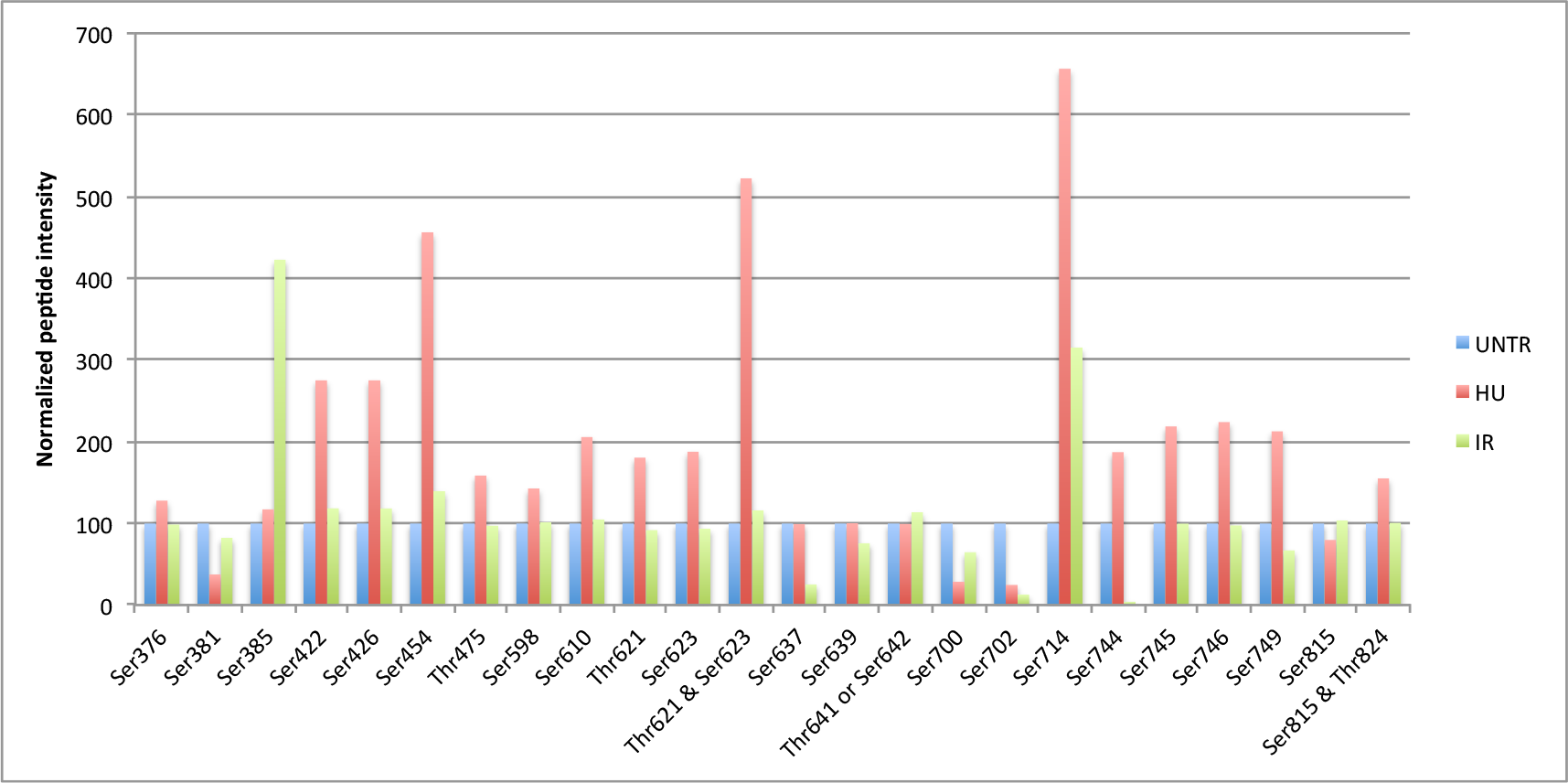
Identification of phosphorylation sites in Strep-EXO1. Graphic representation of intensity signal for the identified phospho-peptides (see Table 2) after normalization to the signal of the same phospho-peptides in the untreated condition.

### EXO1 gene network

To categorize pathway and cellular processes in which EXO1-interacting proteins have an annotated function, we performed gene network analysis of EXO1 binding partners using the literature-based ResNet Mammalian 11.0 Database in Pathway Studio 11.0 [35]. The resulting interaction network displayed connection (i.e., direct interaction) of EXO1 with various cellular pathways, such as RNA and DNA processing, DNA damage response and cell cycle control (Fig. 7). Beside the expected connections with established pathways of DNA resection and remodeling, interaction with RNA processing factors enriches the so far known links of EXO1 with the RNA world, consisting in the ability to phenotypically complement deficiency of Rad27 (the yeast homolog of human FEN1) and displaying RNAse H activity *in vitro* [20]. Our data suggest that EXO1 may be component of mRNA biogenesis complexes that prevent DNA damage generated by aberrant RNA-DNA structures [43, 44].

**Figure 7.**
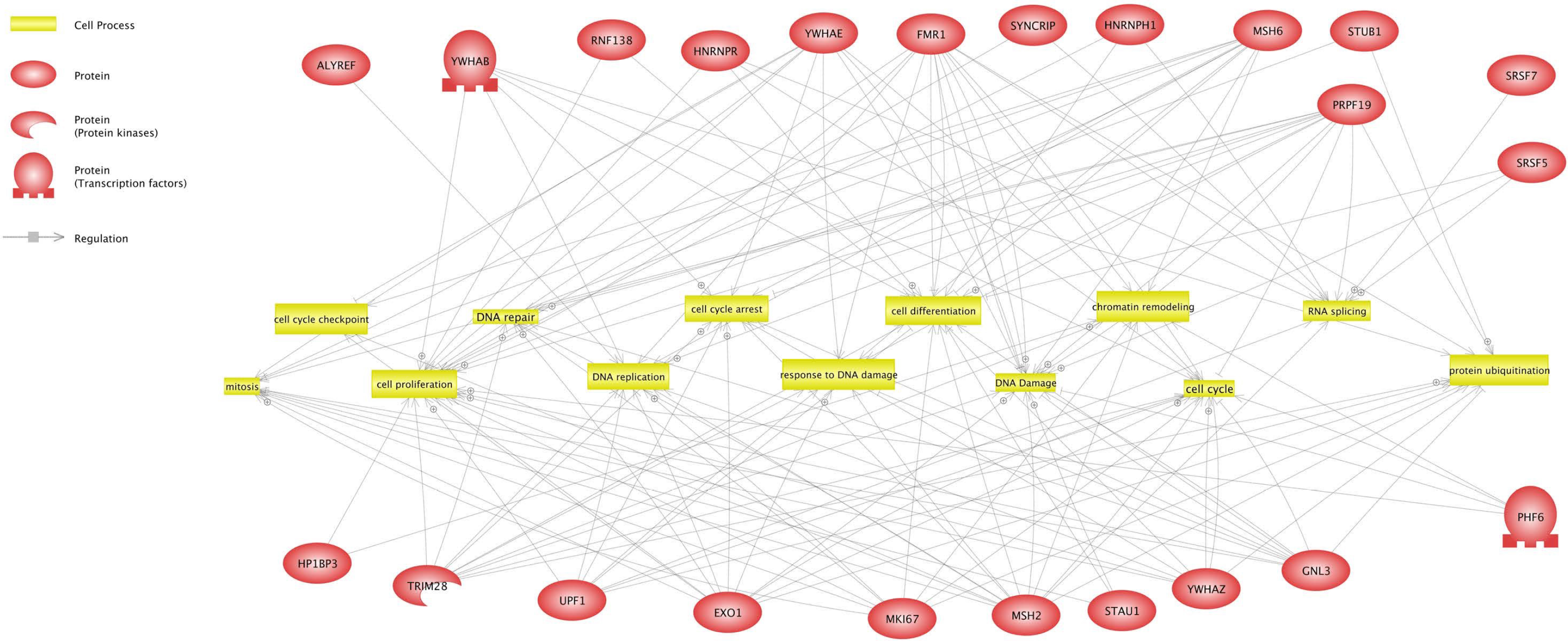
Gene network plot for EXO1 interacting proteins. Gene network analysis of the shortest connections to cellular processes for 24 proteins that interact with EXO1, as obtained by global literature analysis using Pathway Studio 11.0 (Elsevier Amsterdam, Netherlands). Cellular processes are colored in yellow and EXO1 interacting proteins in red. Arrows indicate regulation with the symbol ⊕ representing positive regulation. Bar-headed lines indicate binding.

## DISCUSSION

Dysfunction of the machinery that signals DNA damage and/or addresses DNA repair is associated with cancer development and resistance to therapy [10, 45], providing a direct demonstration of the link between genome instability and cancer [46]. DSBs, are among the most dangerous lesions that may occur to DNA and are generated by internal metabolic processes [3–7] or result from the action of external agents, such as ionizing radiation and mutagenic chemicals [47, 48]. Intense effort is currently being devoted to the identification of proteins and pathways involved in the recognition and repair of various forms of DNA damage to reach a better understanding of the molecular mechanism of action of chemotherapeutics currently deployed in the clinic and, based on the knowledge gained on DDR pathways, to develop more efficient drugs that address the lack of specificity and side-effects of current chemotherapeutics. To allow reaching this second objective, DDR pathway components, their mechanism of action and partner interacting proteins must be identified. In the present work, we focused on the identification of phosphorylation sites and interaction partners of EXO1, an essential component of error-free DNA repair pathways.

To investigate the EXO1 interactome we used a Strep-tagged form of EXO1 and took advantage of affinity-based protein enrichment (Fig. 1). Technically, this approach presents two advantages: (i) it allows eliminating the robust signal of the antibody heavy and light chains that often hampers detection of low-abundance partners of the protein of interest; and (ii) it facilitates identification of weakly interacting proteins by virtue of the mild conditions used for elution of the bait and its partners. Analysis of EXO1 interacting proteins revealed an unexpected network of interaction with proteins involved in RNA metabolism (Table 1 and Fig. 7) possibly suggesting participation of EXO1 in pathways addressing the resolution of RNA-DNA hybrids that threaten genome stability [49, 50]. Other interesting hits were the PHD-finger protein 6 and FMR1, two proteins associated with X-linked disorders, namely the Boerjeson-Forssman-Lehmann syndrome (BFLS) and the Fragile X syndrome / Fragile X tremor-ataxia syndrome (FRAX / FXTAS), respectively. These disorders are characterized by moderate to severe mental retardation and growth defects among other clinical signatures, reminiscent of conditions observed in patients carrying mutations of DNA repair genes [51]. Published evidence indicates that PHF6 deficiency results in DNA damage, which is particularly evident at the ribosomal DNA (rDNA) locus [52], and that FMR1 participates in the DDR thanks to its ability to bind chromatin and regulate the DNA damage machinery [53]. Regarding the possible role of EXO1 in these two syndromes, both FRAX and FXTAS are characterized by abnormal amplification and expansion of CGG repeats [54, 55], suggesting that interaction – and possibly control - of PHF6 or FRM1 on EXO1 as component of the DNA mismatch repair machinery might play an important role in the regulation of repeat expansion [56].

Characterization of a selected subset of EXO1 interacting proteins by knock-down of their expression in cells undergoing DNA damage revealed an important reduction of the DNA damage response (DDR) in all cases examined (Fig. 3 and S2). Among these proteins, the RRP5-homologue and NFκB-interacting protein PDCD11/ALG-4 attracted our attention due to its roles in apoptosis [39], in the generation of mature 18S rRNA [40] and, particularly, because of the frequent aberrations (single nucleotide variations, SNV, and copy number variations, CNV) observed in cutaneous T-cell lymphoma, leading to the proposal of PDCD11 as putative driver gene for this type of pathology [41]. We obtained evidence that PDCD11 depletion rendered cells more resistant to camptothecin or bleomycin (Fig. 4), possibly through a mechanism that results in a decreased response to DNA damage (Fig. 5). In line with our findings, yeast RRP5 was reported to physically interact with Rad27, a 5’ to 3’ exonuclease and 5’ flap endonuclease that is required for Okazaki fragment processing [57], participates in long-patch DNA base-excision repair [58] as well as ribonucleotide excision repair [59] and, like *EXO1*, is member of the *S. pombe RAD2/FEN1* family.

Isolation of EXO1-Strep from HEK293T cells using the StrepTactin system yielded an amount of protein (Fig. 1) sufficient to cover 86% of EXO1 sequence by mass spectrometry. Hence, post-translational modifications such as phosphorylation could be reliably scored (Table 2). Among the sites that we have previously described in EXO1 [25], S422, S454 and S714 were the most prominent residues displaying increased phosphorylation in response to replication stress, with S426 and the doublet T621/S623 being novel sites undergoing similar control (Fig. 6). The newly identified S385 appeared to be the more responsive to ionizing radiation than replication stress (Fig. 6). S714 is an S/T-Q type site, targeted by ATR [25] and to a lesser extent ATM [25, 26], whereas S454 displays the features for recognition by checkpoint kinases (Φ-X_1_-K/R-X_2_-pS/pT, where Φ is any hydrophobic amino acid). S385, S422 and S426 are suitable sites for CK2 (pS/pT-X_0-2_-D/E_1-5_), whereas T621/S623 are typically Pro-directed kinase sites (pS/pT-P). Hence, these data indicate that in addition to classic DDR kinases, also CK2 and possibly stress-activated MAPK members participate in the control of EXO1 in response to genotoxic damage. The majority of the remaining sites, phosphorylation of which did not substantially change in response to DNA damage, could be assigned to the AGC and the CMGC subfamilies of protein kinases or to CK1. Among the residues targeted by Pro-directed kinase are cell cycle regulated sites, the phosphorylation of which was shown to depend on CDK2 in S- and G2-phase and to peak in prometaphase-arrested cells when CDK1 activity is maximal [28].

The identification of novel EXO1 interacting proteins presented here, such as the causative agents of X-linked syndromes, which have been previously linked to DDR, or PDCD11, which represents a novel entry, warrants future studies on the molecular mechanism by which these proteins specifically affect EXO1 function and, in general, participate in cellular responses to genotoxic damage. The additional comprehensive identification of EXO1 phosphorylation sites adds another layer of complexity to the mechanism that control EXO1 localization and function at sites of damage.

## ADDITIONAL INFORMATION

Annotated, mass labeled spectra and all data for the analysis described in this study are available at ProteomeXchange with the following identifiers:

Title: Exonuclease-1 interactome and phosphorylation sites
Accession: PXD002780
Project webpage: https://www.ebi.ac.uk/pride/archive/projects/PXD002780

## AUTHORS CONTRIBUTION

W.E.: Contributed to conceive the study and performed experiments in Fig. 1-6-7, Table 1-2 and analyzed the data;

D.H.: Performed mass spectrometry and analyzed the data;

C.K.: Performed experiments in Fig. 3-4-S2 and contributed to experiments in Fig. 2-5-S3;

C.G.: Analyzed the data in Table 1;

SF: Conceived the study, acquired funding, designed and performed experiments in Fig. 2-5-S3, analyzed the data and wrote the manuscript.

## ACKNOWLEDGMENTS

We are indebted to F. Mhamedi for invaluable technical support. We would like to thank members of SF laboratory for helpful suggestions.

This work was supported by grants obtained from Swiss National Science Foundation (SNSF 31003A-144009), Promedica-Stiftung, Hartmann-Müller-Stiftung, Swiss Foundation for Fight Against Cancer (SKB-355), Hermann-Stiftung, Huggenberger-Bischoff-Stiftung and University of Zurich Research Funds (SwF-783).

## STATEMENT

We declare that all methods were carried out in accordance with relevant guidelines and regulations. We also confirm that experimental protocols were approved by the University of Zurich.

## TABLES

**Table 1** – List of 385 EXO1 interacting proteins based on exclusive spectra counts, with indication of gene and protein name. The CRAPome cut-off value was 206. Entries confirmed by co-immunoprecipitation studies (Fig. 2) are indicated with an asterisk. Entries analyzed in RNA-interference studies are indicated with a double asterisk.

**Table 2** – List of the all peptides analyzed for the presence of post-translational modifications. Phosphorylated amino acids present in phospho-peptides among the identified peptides are indicated in red (pS = phospho-Serine; pT = phospho-Threonine). Only phospho-peptides identified at least twice were considered. Phospho-amino acids in italics indicate dubious assignment of the phosphorylation site.

---

**Supplementary Figure S1 – Verification of EXO1 interacting proteins**

HEK-293T cells were transfected with an empty plasmid or a plasmid expressing EXO-Strep and the deriving WCE (2 mg) submitted to pull down. Proteins were resolved by SDS-PAGE and, upon transferring to PVDF, the indicated proteins were revealed with specific antibodies. Inputs correspond to 50 μg WCE.

**Supplementary Figure S2 – Effect of EXO1 interacting proteins on the DDR**

U2-OS cells transfected and treated as described in the legend to Figure 3 were visualized by fluorescence microscopy. Sample images for the quantification presented in Figure 3 are shown.

**Supplementary Figure S3 – Characterization of siRNA oligonucleotides to PDCD11**

A. U2-OS cells were transfected with two distinct siRNA oligonucleotides to PDCD11 used alone or in combination. Depletion of PDCD11 expression was verified on whole cell extracts 48h upon transfection. KAP1 was used as loading control.

B. Cell cycle effect of PDCD11 depletion with two distinct siRNA oligonucleotides used alone or in combination. Acquired events are shown as dot plots. Gated regions represent G1, S and G2-M cells, respectively.

C. The same data presented in B. were visualized in histograms.

**Supplementary Table 1.**
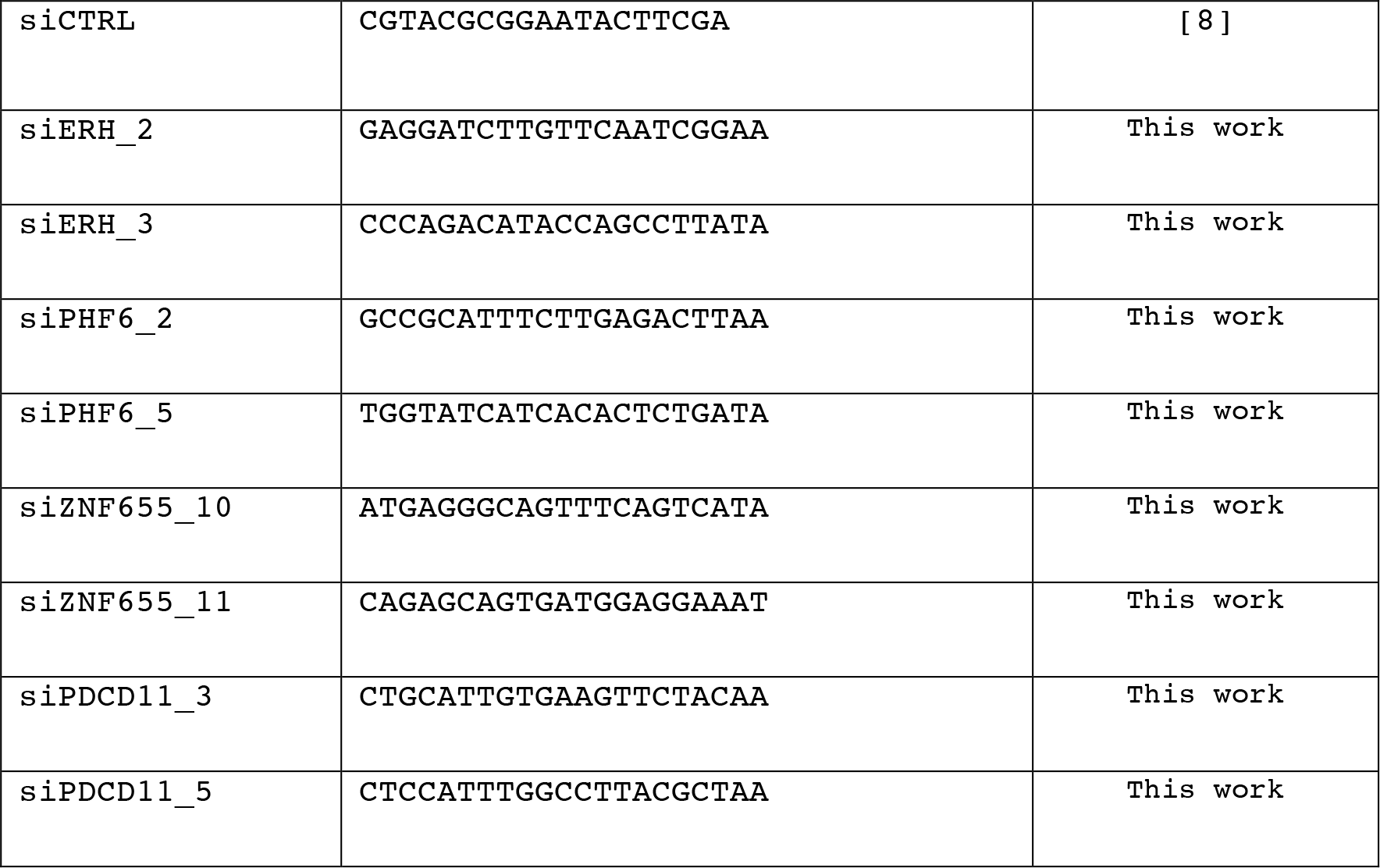
List and references for the oligonucleotides used in RNA interference experiments.

