## Supplementary figures and images for "The Human Exonuclease-1 Interactome And Phosphorylation Sites"

### Supplementary Figures

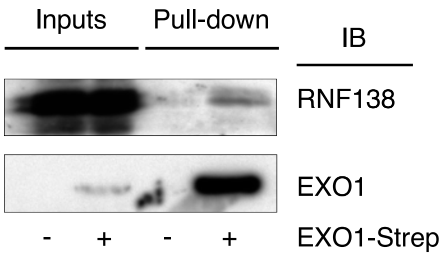


Eid et al. Fig. S1

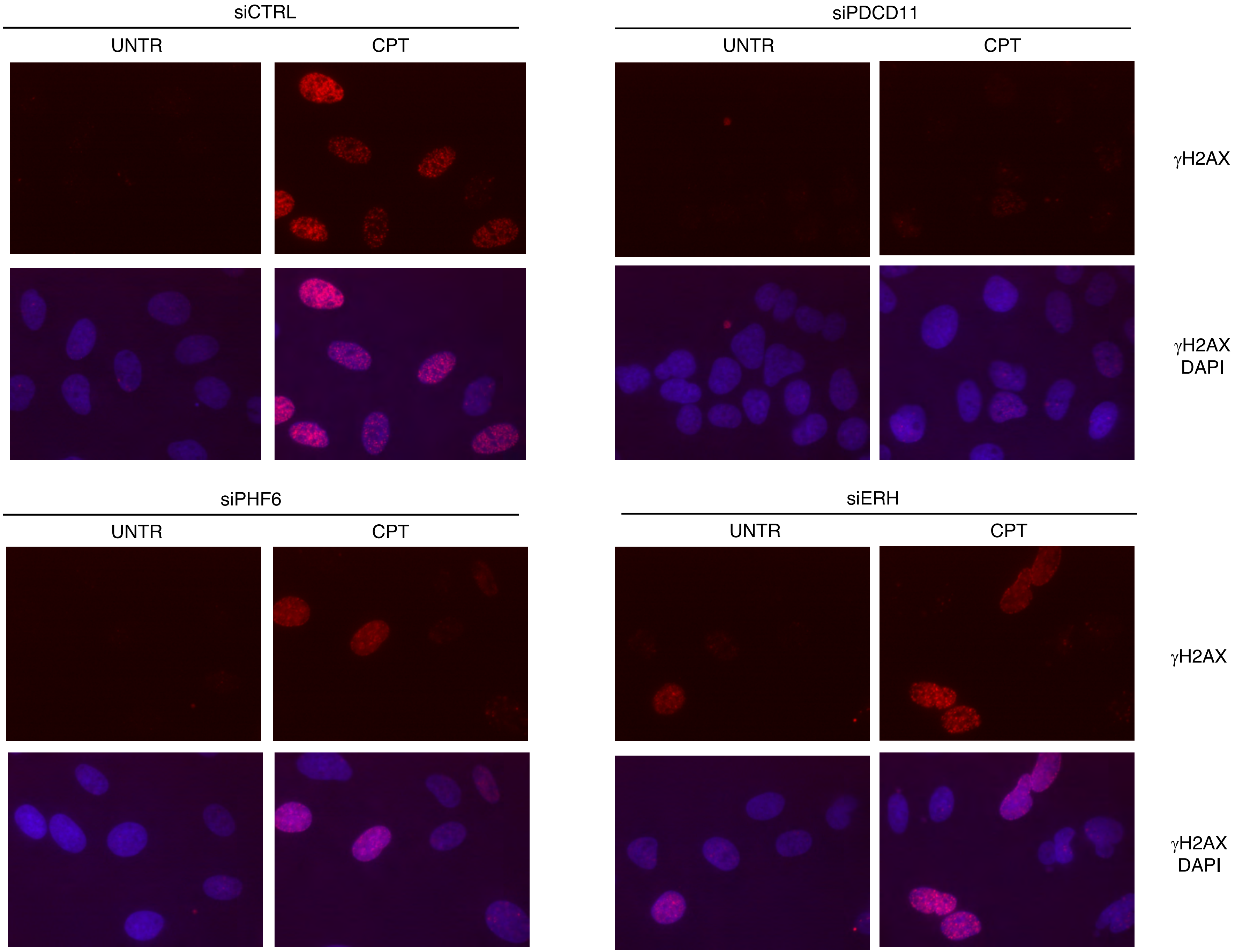


Eid et al. Fig. S2

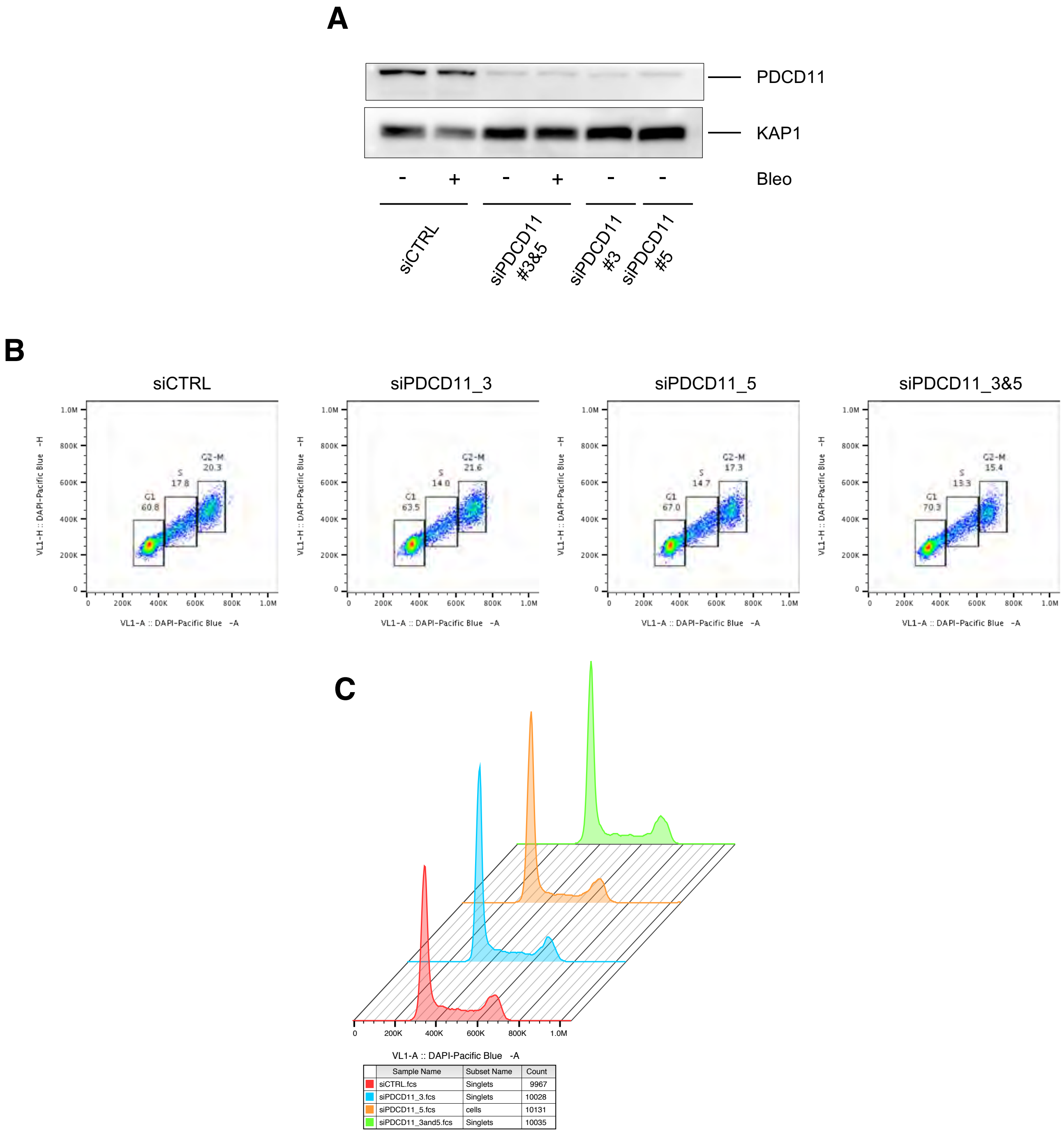


Eid et al. Fig. S3
